# Exploratory movement analysis and report building with R package stmove

**DOI:** 10.1101/758987

**Authors:** Dana P. Seidel, Eric R. Dougherty, Wayne M. Getz

**Affiliations:** Environmental Science, Policy & Management, University of California, Berkeley, USA; School of Mathematical Sciences, University of KwaZulu-Natal, Durban, South Africa

## Abstract

**Background:** As GPS tags and data loggers have become lighter, cheaper, and longer-lasting, there has been a growing influx of data on animal movement. Simultaneously, methods of analyses and software to apply such methods to movement data have expanded dramatically. Even so, for many interdisciplinary researchers and managers without familiarity with the field of movement ecology and the open-source tools that have been developed, the analysis of movement data has remained an overwhelming challenge.

**Description:** Here we present `stmove`, an R package designed to take individual relocation data and generate a visually rich report containing a set of preliminary results that ecologists and managers can use to guide further exploration of their data. Not only does this package make report building and exploratory data analysis (EDA) simple for users who may not be familiar with the extent of available analytical tools, but it sets forth a framework of best practice analyses, which offers a common starting point for the interpretation of terrestrial movement data.

**Results:** Using data from African elephants (*Loxodonta africana*) collected in southern Africa, we demonstrate `stmove`’s report building function through the main analyses included: path visualization, primary statistic calculation, summary in space and time, and space-use construction.

**Conclusions:** The `stmove` package provides consistency and increased accessibility to managers and researchers who are interested in movement analysis but who may be unfamiliar with the full scope of movement packages and analytical tools. If widely adopted, the package will promote comparability of results across movement ecology studies.

## 1 Background

With the increased accessibility of GPS data and expanded computing power for analyzing such data, a concomitant open-source software expansion has occurred for exploring spatiotemporal structure in movement trajectories. With this expansion of data and tools, two vexing problems remain for researchers and managers: 1.) lack of a unifying movement-pathway framework that would facilitate comparisons across studies; and, 2.) lack of software-package accessibility (e.g., using R or Matlab) to those not steeped in movement ecology or those lacking proficiency in command line program implementation.

These two problems are exacerbated by the sheer volume of available tools, which has resulted in an overwhelming landscape of analytical options. This is especially challenging for researchers or managers who fortuitously get access to GPS relocation data, but have little or no experience in analyzing movement trajectories. R has emerged as an open source programming platform of choice for movement ecologists, primarily because of the large number of statistical and data manipulation packages that have become available to aid researchers in conducting even the most obscure domain-specific analyses. In a recent review of R packages solely dedicated to movement or tracking analyses, 57 individual packages were listed, with only 12 having good to excellent documentation [1]. It is also clear that, while these open-source, programming languages—with R being a primary example—have made it possible for scientists to carry out an ever-growing list of new and developing analyses, these packages constitute a variety of different, and not all together compatible, methods and data structures. Thus, this burgeoning cornucopia of tools can be as much a stumbling block as a godsend for many researchers and managers: even with access to a trove of GPS movement data, they may not have the time or expertise to assimilate which of the available toolkits is most appropriate for their analytical needs.

Building on the work of previous open source contributors, our `stmove` R package alleviates the first of these problems—i.e., lack of a coherent framework—by setting forth a standard set of first-pass exploratory analytical methods that should be performed before undertaking more specific or targeted movement analyses. Further, it mitigates the barrier-to-use problem by conveniently gathering in one place a disparate set of compatible tools and methodologies and providing a single command (and optionally interactive) infrastructure for automated report building that can run analyses and compile results into a digestible, visually rich report. From this report, researchers and managers can then distill key insights needed to sharpen current interpretation and subsequent exploration of their movement data. Of course, our package cannot streamline research to the point where no additional analyses are needed: many questions require deep methods that are too sophisticated to be included in a general entry-level R package. Instead, our package seeks to make it much easier to carry out a first cut analysis, using a standard set of methods. We propose that such analyses, as detailed in the implementation section, should to be undertaken before more powerful methods, required to address complex questions, are applied.

## 2 Implementation

R is an open source and system-agnostic language [2], with a growing user base in the ecology and environmental science communities. Our `stmove` package can be used on any computer with R installed regardless of operating system (e.g., MacOS, Linux, or Windows). Alternatively, `stmove` may be used within a web browser when combined with external cloud computing services such as RStudio Cloud. The primary goal of `stmove` is to preform a standard set of exploratory data analyses and return a preliminary report with visualizations, interpretation aids, and suggested next steps to empower managers or researchers new to movement analysis. In addition, it aims to give all users a simple work-flow of best practice analyses from which to begin any movement study. Although all analyses are standalone and can be preformed separately on individual trajectory data, the primary advantage of this package is its report building function that conducts multiple analyses and provides a PDF report for each inputted set of movement trajectory data. This report includes 4 central components: general distributions (step-size and turning angles [3]), interval statistics (means, variances, auto and cross-correlations, and plots of running averages of these), wavelet analyses [4, 5], and home-range constructions [6, 7]. Additionally,`stmove` functions to summarize multiple trajectories in space and time. Each of the packages main components and their underlying methods are discussed below. Software design choices have been made to further simplify the use of `stmove`, including sensible defaults and an interactive HTML add-in to help guide report building for users implementing `stmove` within the popular RStudio IDE (integrated development environment).

### Individual analysis

The goal of `stmove` is to make more accessible the standard spatiotemporal approaches to analyzing and interpreting movement data before implementing project-specific approaches to deconstructing movement trajectories. Application of `stmove` requires a clean, regular, GPS time series of relocation data consisting of a sequence of *T* + 1 points (*x*_*t*_, *y*_*t*_, *t*) where *t* = 0, 1, 2, 3, *…, T*, and all missing points have been interpolated and filled in. Additionally, locations (*x*_*t*_, *y*_*t*_) are expected to be in a projected coordinate system—the unit of measure in the coordinate system determining the unit of measure for calculating the step-size (*s*_*t*_) and turning-angle (*θ*_*t*_) time series of lengths *T* and *T -* 1 respectively [3], using the equations

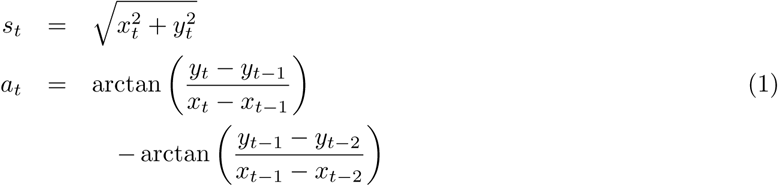

All analyses are intended to be performed on a single individual’s trajectory to provide insight on individual movement patterns; see our population analysis section to review the analyses available for multi-animal datasets supported by `stmove`.

### Regularization and Interpolation

Ensuring that relocation data are complete and regular—that is, the data collection frequency must be fixed and no values should be missing—can represent a sufficiently challenging hurdle to movement analyses that it may be the reason why GPS data often go unanalysed. Even for telemetry data collected at a set fixed rate, relocation timestamps can be imprecise, recorded within a few seconds or even minutes of the scheduled time, due to lags in satellite connections, and fixes may be missed altogether for a variety of reasons.

To help researchers easily regularize near-regular data, the popular `adehabitatLT` R package [8] has two functions ‘setT0’ and ‘setNA’ which `stmove` has wrapped together in a new function called ‘regularize’ to help users easily regularize their data. Given a reference date and time and an expected fix rate, ‘regularize’ will round fix times to the nearest fix (within a set tolerance) and insert NAs (formal missing value notation) for all missing values to build a regular time series. Because further analyses require complete trajectories, `stmove` then provides the function ‘kalman’ for interpolation of missing values.

The ‘kalman’ function uses Kalman smoothing to interpolate missing values in the *x* and *y* coordinates of a trajectory. The Kalman smoothing approach is a state-space model that uses all available observations (rather than just the past and current observations, as in the case of Kalman filtering) to derive the covariance structure of the data and predict the current state. This function, based upon the ‘na.kalman’, ‘StructTS’, and ‘auto.arima’ functions of the `imputeTS` package [9], selects the best fit structural (linear, Gaussian) time-series model for each univariate time series in turn—i.e., treating the series *x*_*t*_ and *y*_*t*_ separately— and then applies Kalman smoothing (based upon the chosen best fit models) to iteratively interpolate any missing values. Since our intention is to provide a rapid, flexible way to interpolate missing points for the purposes of exploratory data analysis, `stmove`’s ‘kalman’ function is optimized for rapid estimation rather then the most accurate possible interpolations. For this reason `stmove`’s ‘kalman’ implementation reports the ratio of interpolated to empirical points and issues a warning to users when interpolating more than 5% of a trajectory’s total points. Because errors associated with interpolation degrade the accuracy of ensuing visualizations and analyses knowledge of this ratio forms part of an assessment (albeit informal) of the reliability of the results obtained. If users are seeking to analyze trajectories with large gaps between otherwise consistent fix intervals, they are encouraged to break trajectories up into sub-individual trajectories or choose the largest continuous section of sampling for subsequent analysis rather than interpolating over large gaps. For intentionally gappy or opportunistically gathered telemetry data (e.g., from marine mammals when surfacing, or from older or failing satellite collars), more rigorous models for interpolation may need to be considered. These methods, however, are outside the scope of the `stmove` package and are discussed elsewhere [10, 11, 12].

### Visualizations and Distributions

Before undertaking any exploratory data analysis, it is generally helpful to visualize the data. In the case of movement data, visualization of a trajectory can show outliers, recursions [13, 14] and syndromic movement behaviors [15], and the individual’s general space use pattern. Implemented using R’s `ggplot2` package [16], the first plot returned by `stmove`’s ‘build report’ function is a simple scatter plot of the (*x, y*) coordinates of all locations defining a given trajectory (e.g., see Fig. 4). The next plots in the report are histograms of step size *s*_*t*_ and turning angle *a*_*t*_. These are obtained by `stmove` converting a user-provided dataframe of relocations to an ‘ltraj’ object (an object class defined by the popular `adehabitatLT` package [8]) and then calculating the step-size and turning-angle time series using Equations 1 (e.g., Fig. 5). `stmove` then plots step-size and turn-angle histograms using `ggplot2` (e.g., see Fig. 5; [16]). From these plots depicting the frequency of different step sizes or turning angles, users can begin to identify outliers and perhaps detect behavioral modes or directional biases in the trajectory.

**Figure 1:**
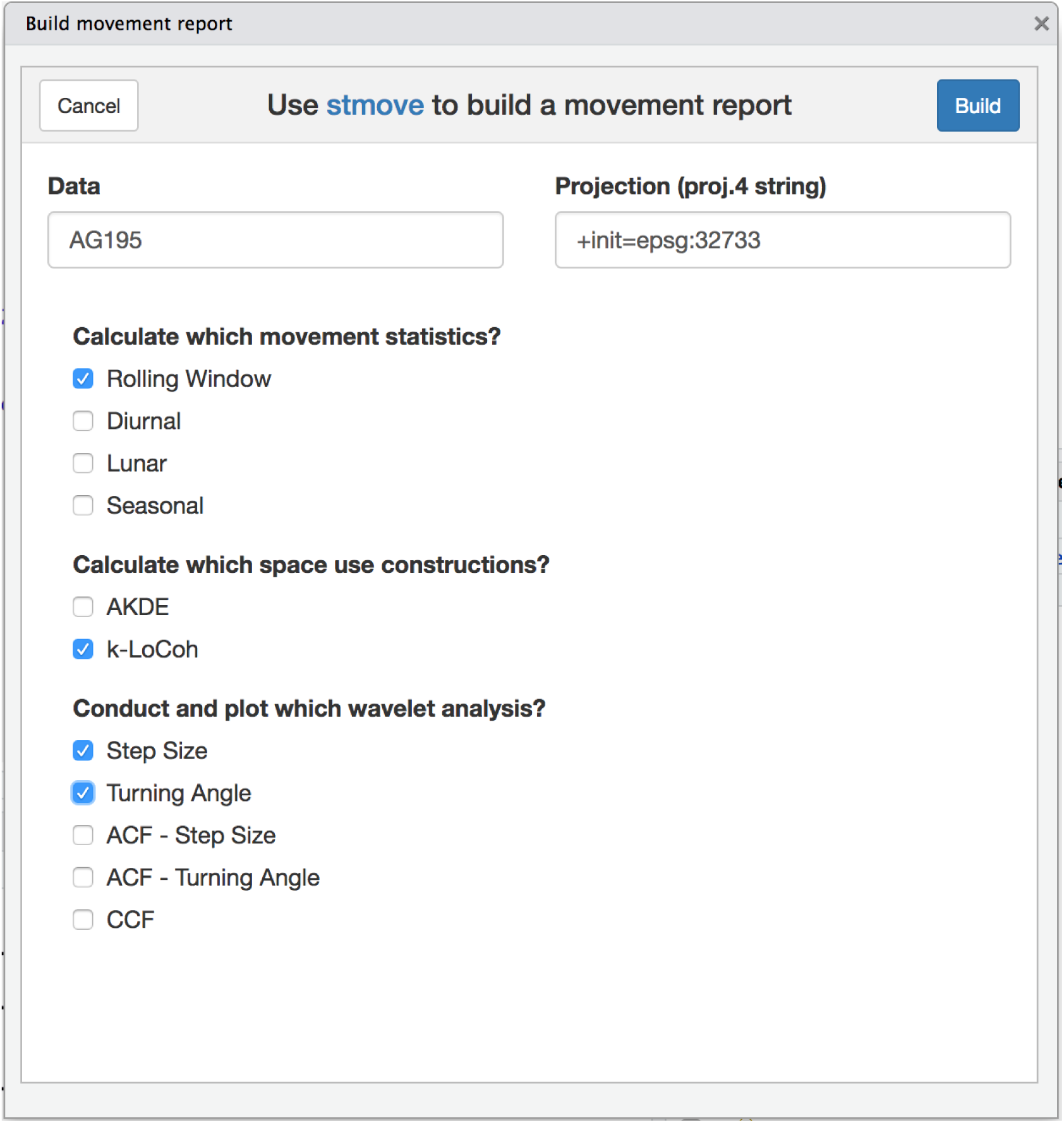
Rstudio AddIn Menu. A screenshot of the `stmove` Report Builder add-in. This interactive menu guides users of the package within RStudio through the customization of their movement reports.

**Figure 2:**
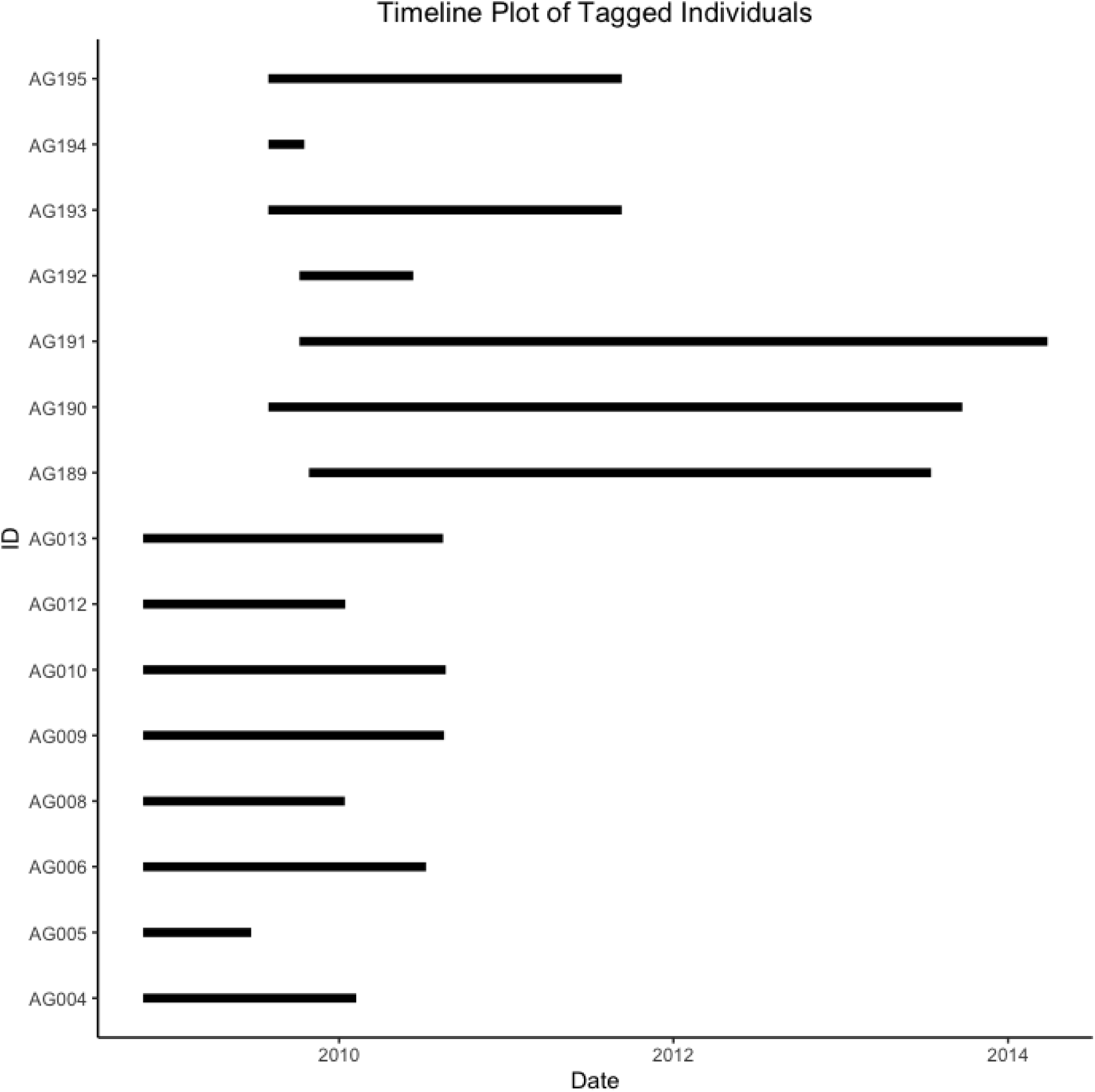
Sampling Timeline Plot. A segment graph demonstrating the sampling period for each individual in the elephant dataset. This is the output of `stmove`’s ‘plot timeline’ function and is called by ‘build report’ when given a data frame including multiple ids.

**Figure 3:**
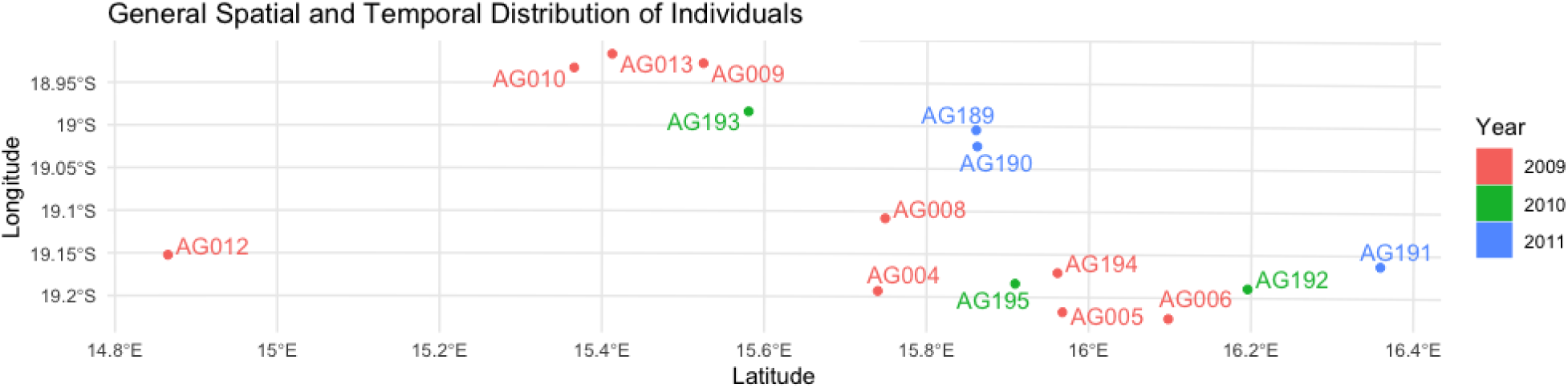
Spatial Temporal Distribution Plot. A map demonstrating the mean x and y locations of 15 individuals included in the elephant dataset.

**Figure 4:**
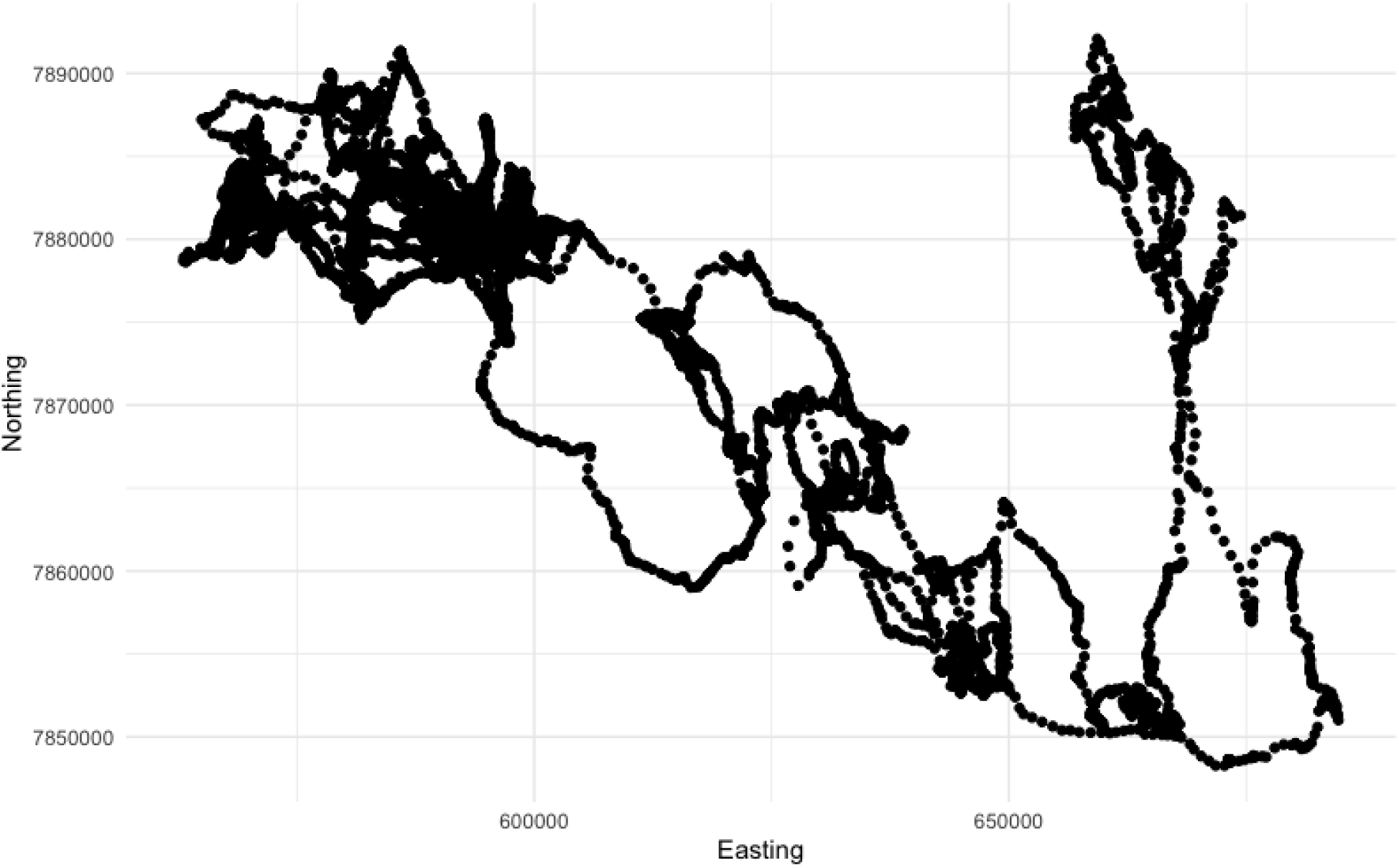
Simple Coordinate Plot. A simple *x*-*y* plot of coordinates along AG195’s trajectory.

**Figure 5:**
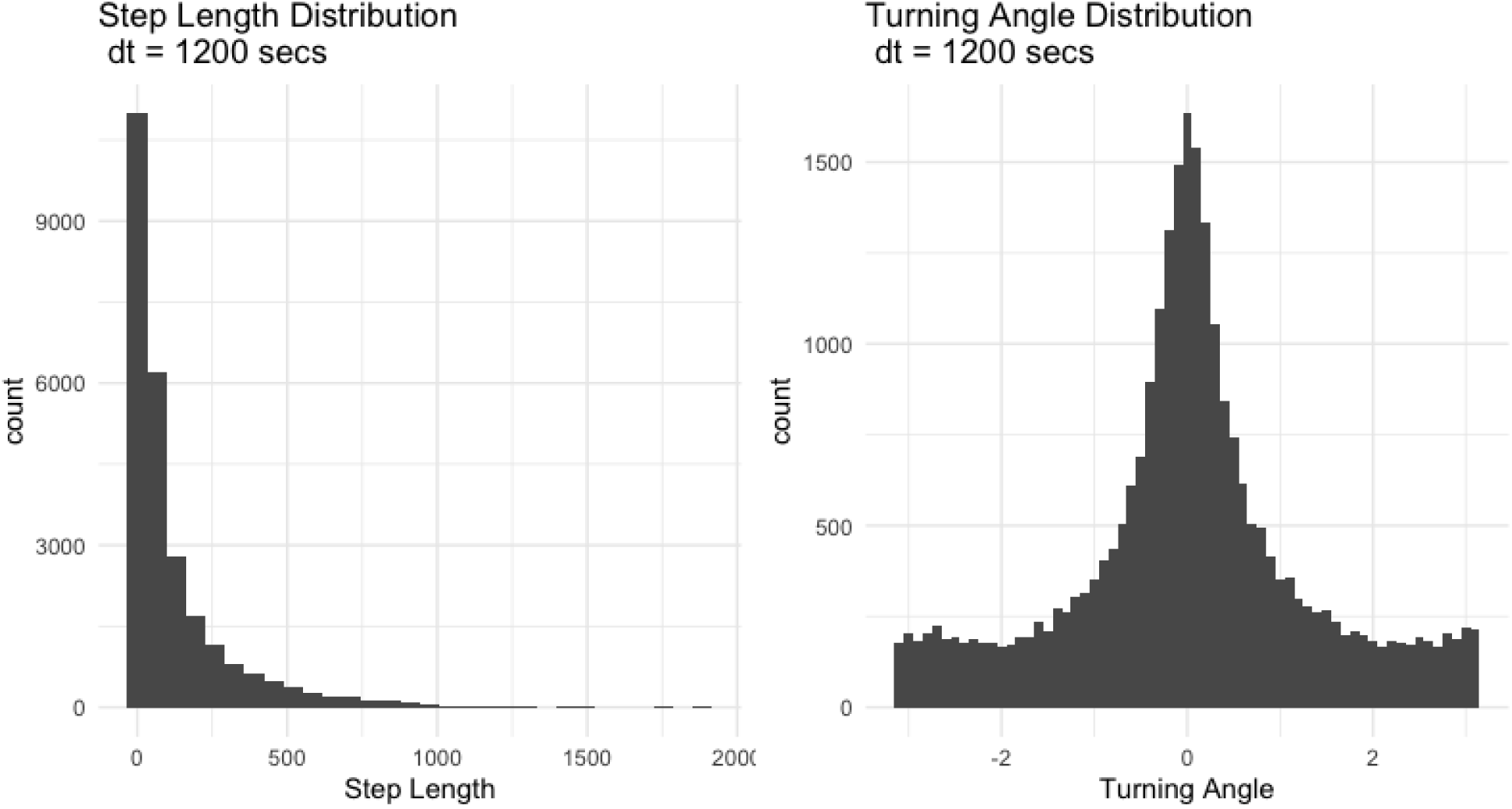
Step Size and Turning Angle Distributions. `stmove` reports include two histograms visualizing the distributions of primary movement metrics step size and relative turning angle.

#### Rolling Statistics

Our next step is to calculate the primary times series statistics using rolling (sliding) windows *W* (*t, w*) of a given fixed length *w* and starting at time *t*. Within windows *W* (*t, w*) for *t* = 0, *…, T w*, `stmove`’s ‘rolling stats’ function computes running step-size and turning-angle means, as well as single-lag autocorrelations and cross-correlations of the *s*_*t*_and *a*_*t*_time series. These rolling windows are implemented using `RcppRoll`’s ‘roll meanr’ and ‘roll sdr’, and `TTR`’s ‘RunCor’ functions [17, 18], all statistics are calculated with a right-aligned window. If fix rate is sub-hourly, the rolling window defaults to three hours, if the fix-rate is one hour or more, the window is increased to six hours. These defaults can be overridden with the optional ‘n roll’ parameter allowing users to specify how many fixes they wish to “roll” over when calculating statistics. This argument is particularly powerful if users are investigating a trajectory with a large fix rate (i.e., *>* 3 hrs), for which the default behavior will not provide especially informative results. Rolling statistics are often used as inputs to more advanced types of movement analyses [19, 20]. These rolling window plots offer users insights into behavioral patterns that may relate to the identification of different modes of activity (e.g., using break-point analyses [21, 22]). In addition, we note that auto- and cross-correlations are transformations of primary movement metrics that can estimate persistence in either direction (acf ang) or distance/speed (acf dist).

#### Interval Statistics

While rolling statistics can smooth patterns through time, interval statistics are a preliminary means to identify patterns across discrete, biologically meaningful periods of time. `stmove`’s ‘interval stats’ function can be used to calculate the mean and variance of a trajectory’s primary movement statistics across three intervals of interest: diurnal, lunar, and seasonal. Diurnal analysis summarizes these statistics for 12 hour windows representing pre-noon (0-12) and post-noon (12-24) hours, as determined by the time zone associated with user-inputted data. Lunar interval analysis relies on the `lunar` package [23], automatically dividing a given trajectory according to periods within the lunar cycle, full-waning, and new-waxing intervals according to date. Seasonal interval analysis is customizable with `stmove`: ‘interval stats’ recognizes an optional ‘seas’ argument by which users provide a character vector of season start dates. In this way, users are allowed to specify custom seasons over which to calculate the interval statistics. Which interval statistic is appropriate may depend upon the length of a users trajectory and/or the biology of the tracked animal. As such, when building a report, the user can specify which interval statistics they would like to include using the ‘stats’ argument.

#### Wavelet transform and visualizations

Many factors influencing movement are cyclic with periods that are linked to ecological relevant frequencies (e.g., regular resource gathering trips, migration, or certain social and reproductive behaviors). Fourier and wavelet transformation methods are useful analyses to examine the cyclical nature of animal movement and behavior [4, 5]. Especially as a part of exploratory analyses, these time-frequency methods are useful for understanding dynamic movement responses to physiological, ecological, climatic, and landscape factors [4, 5]. After `stmove` has calculated basic path distributions and statistics, it implements a wavelets analysis on user-selected time series, using Morelet filters, by importing functions from an existing open source R package, `dplR` [24, 25]. The user-selected time series are step sizes, turning angles, autocorrelation coefficients of both coefficients and their cross-correlation coefficient. It then produces a plot of the local wavelet power spectrum that users can then use to visually identify any possible periodic components (e.g., see Fig. 7).

**Figure 6:**
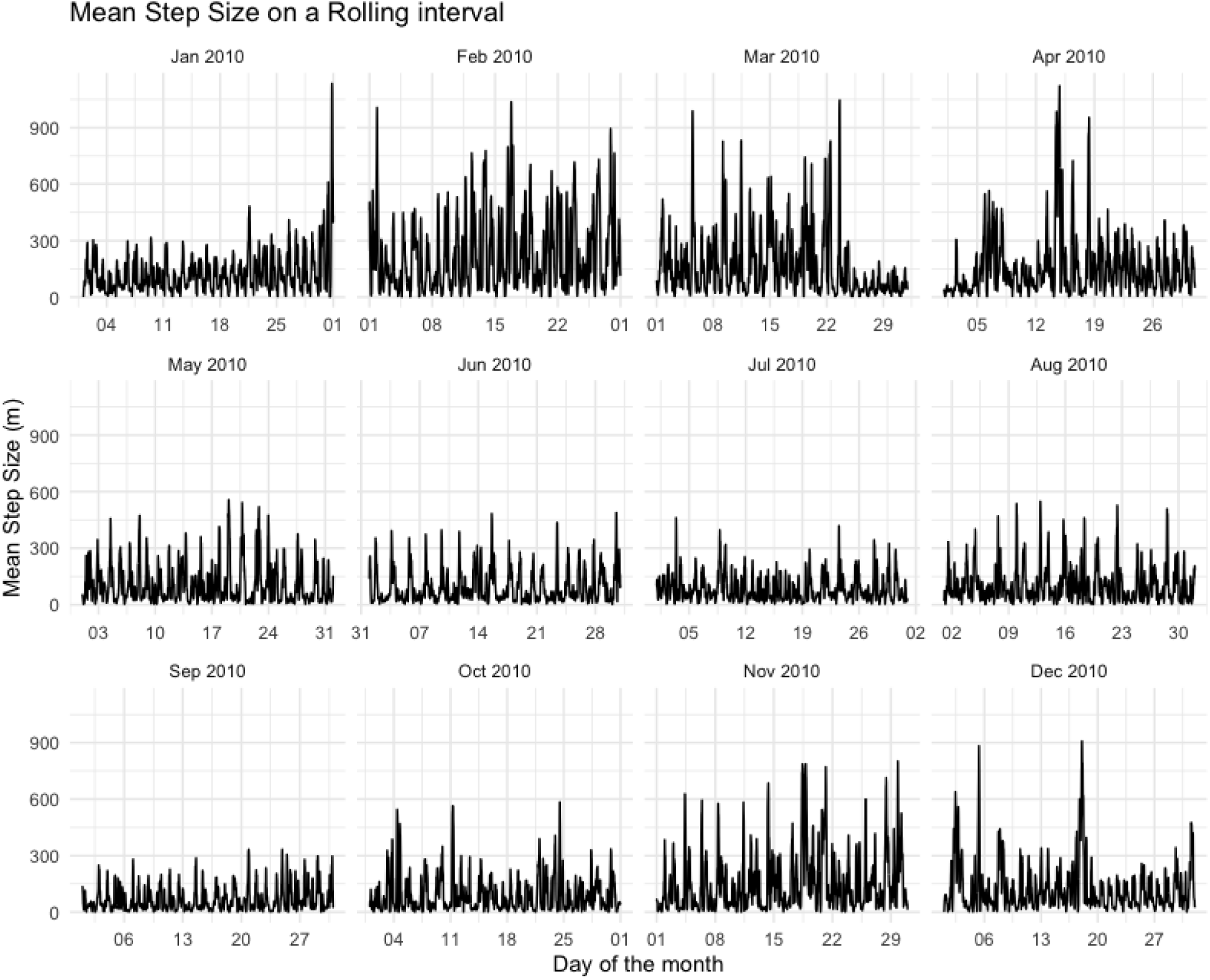
Rolling Step Size. A faceted plot of step size averaged using a rolling window of 3 hours using `stmove`’s ‘rolling stats’ function. `stmove` reports include plots of rolling means for step length (*s*_*t*_) and turning angle (*θ*_*t*_), as well as rolling autocorrelations of both (*s*_*t*−1_*s*_*t*_ and *θ*_*t*−1_*θ*_*t*_) and a rolling cross-correlation between them (*s*_*t*_*θ*_*t*_)

**Figure 7:**
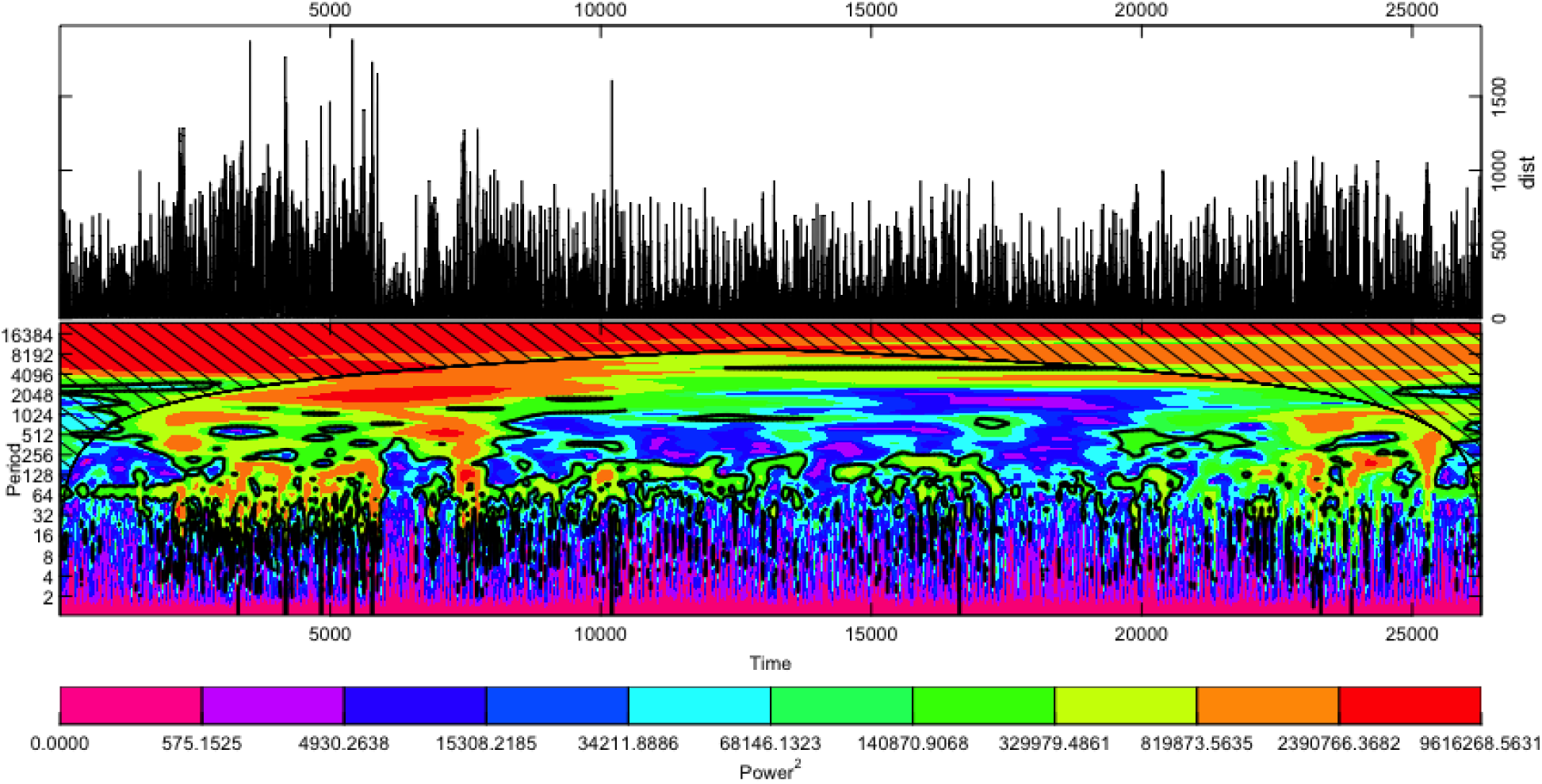
Wavelet plot. In the top half of the plot, the time series used for the wavelet analysis, in this case step length is plotted. In the lower plot, the power spectrum of the Morelet wavelet transform of this statistic is plotted. The lower left axis is the Fourier period corresponding to the wavelet scale of the top right axis. The bottom and top axes display time, represented simply as a time series index. Thus the units of all axes are subject to the underlying fix rate of the trajectory, in this case 20 minutes. The coloured *power*2 contours are added for significance, the thick contour encloses regions of greater than 95% confidence. Cross-hatched regions on either end indicate the “cone of influence,” where interpretation may be impacted by edge effects and should be avoided. These power spectrum plots are one way to investigate underlying periodicity in movement behavior from a trajectory’s primary statistics.

#### Basic space constructions

A crucial step to exploring movement data is understanding what a given trajectory can tell us about higher order space use, notably an animal’s core area or home range [7]. There has been considerable debate regarding the best tools for evaluating landscape level space use, including methods from minimum convex polygons that are conceptually simple and computationally cheap to more complex methods incorporating an animals probability of occurring in a given location [26]. `stmove` incorporates two non-parametric spatial construction methods for users to choose among when estimating 25, 50 and 95% home range isopleths: a Local Convex Hull construction implemented with the `tlocoh` package and an auto-correlated utilization distribution analysis implemented with the ‘akde’ function from the `ctmm` package [6, 27, 28, 29] (e.g., see Fig. 8). Both methods have their particular strengths: the `ctmm` AKDE method (i.e., implemented using the ‘akde’ function) provides a statistically rigorous construction when analysing correlated data under the assumption that movement is an Ornstein-Uhlenbeck process—i.e., a continuous time generalization of an autocorrelated random walk, sometimes with drift added [28, 30]; and, when the latter assumption is not valid (e.g., when the frequency of relocation sampling is at the same or longer time scales for which movement decisions are influenced by environmental factors) `tlocoh` implicitly accounts for vegetation and landscape structures, as well as hard boundaries due to irregular landscape features [6, 31, 32]. In `stmove` we implement *k*-LoCoH with 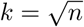(rounded to the nearest integer, where *n* equals the number of relocations in the time series). We also note that the `ctmm` ‘akde’ method produces space-use estimates with confidence intervals that appropriately account for the autocorrelation inherent in movement data. Either method can be implemented using the ‘construct’ function and specifying the method with the ‘type’ argument.

**Figure 8:**
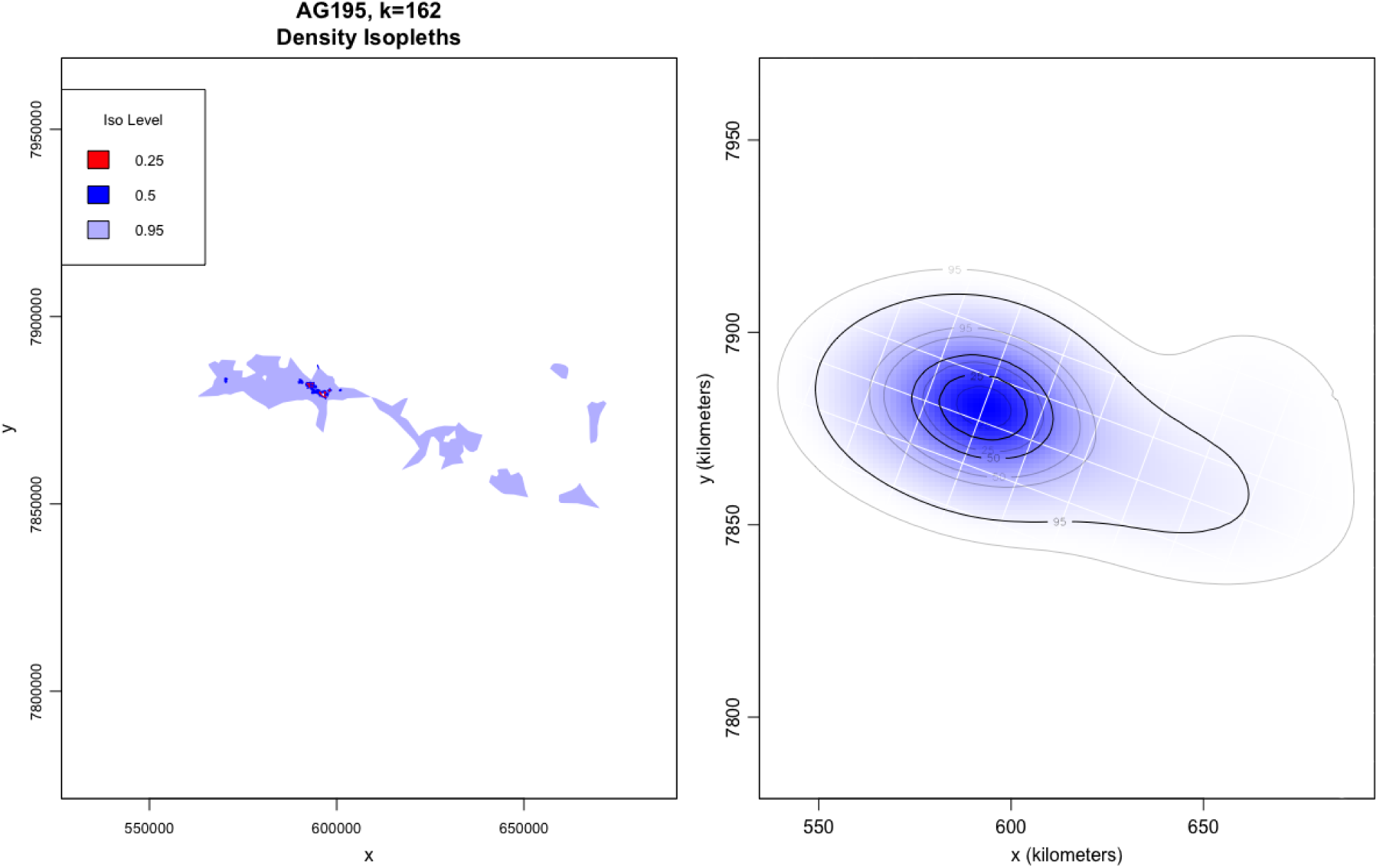
Spatial Constructions. A demonstration of `stmove`’s two spatial construction types, *k*-LoCoH and autocorrelated kernel density estimation. Note the difference in area estimation across the two non-parametric techniques.

### Population analysis

Though the package is designed to build reports for individual trajectories, when provided with a data frame storing multiple trajectories specified by four columns—*x, y*, date, and id—`stmove` is able to create an additional cover sheet showing spatial and temporal overlap of the individuals in the dataset using two important functions: ‘plot_timeline’ and ‘dist_map’. These plots, built using `ggplot2` [16], summarize the spatial and temporal overlap of trajectories. In the first plot, a segment graph produced by ‘plot_ timeline’ is displayed, with lines identifying the duration of time sampled for each individual in the data frame. In the second plot, built by ‘dist_map’, all of the included trajectories are plotted with a single point representing the mean *x* and *y* coordinate over the course of the full path and colored according to the mean year of sampling. These visualizations provide a straightforward summary of the spatial and temporal spread of trajectories within a data set (see Figs.2, 3).

### Report Building

The primary product of this package is ‘build_report’, which produces a PDF report of the results of the analyses described above, when given a clean, regularized trajectory. Beyond the initial three plots discussed in the *Visualizations and Distributions* subsection, which are included in all reports, reports can be customized to include any or all of the analyses by changing the arguments passed to ‘build_report’. ‘build_report’ is implemented using the `rmarkdown` package [33, 34] and parameterized templates that are distributed and installed with the package. The templates use the popular visualization package `ggplot2` [16] to deliver users a report with custom visualizations of their selected analyses. To aid users in customizing their reports, we have implemented an interactive HTML widget that can help guide users of the popular IDE RStudio in building their own movement reports (Fig. 1).

## 3 Illustrative Example

We illustrate the implementation of the `stmove` package using relocation data collected from a population of African Elephants (*Loxodonta africana*) in and around Etosha National Park, Namibia. We generate our `stmove` analyses and report using a data frame containing 15 individual trajectories previously published by Tsalyuk et al. [35]. After initial regularization, each individual trajectory contained between 5633 to 113652 empirical relocations. This unique data set contains individuals tagged from October 2008 through July 2015 for variable-length sampling periods and fixed rates.

### Population Analyses

For data sets with multiple trajectories, the `stmove` population functions provide a powerful summary of our complete data set in a population level “cover page”. The ‘plot_timeline’ function produces Fig. 2, a plot that immediately captures the coverage of and variability in sampling intervals across the population. In a second plot, produced by the ‘dist_plot’ function, Fig. 3, the spatial and temporal distribution of individuals is displayed by plotting for each individual its the mean location 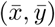from all fixes available. Each mean x-y point is then colored according to the average year of the relocations. These abstractions distill 724,925 points and 15 individuals concisely to communicate the spatial and temporal spread of our population. From these plots we see that we a group of one individual on the eastern end of our space, a group of four in the center top and a string individuals in the eastern bottom half of our space. have at least one group of individuals clustered very close together, perhaps because of habitat constriction, opportunistic sampling, or social structure. This sort of result immediately spurs questions for further analysis, including those concerning the social structure of our population of interest, exemplifying the purpose of solid exploratory data analyses and the `stmove` package.

### Data regularization

To demonstrate the per-individual metrics of the package, we use data from individual AG195, a female elephant collared from July 2009 through September 2011 with fixes taken at consistent but irregular intervals, with points collected in repeating intervals of 1 minute and 19 minutes. Before analysis with `stmove`, regularization was performed using ‘regularize’ and an expected fix rate of 20 minutes – eliminating every other fix in order to standardize the interval to 20 minutes for future analysis (50% observation loss). The complete regularized trajectory consisted of 55,524 relocations but for ease of demonstration and interpretation we will use only relocations from 2010 in the following analyses (*n* = 26277). This regularization procedure was followed by the execution of the ‘kalman’ function to interpolate 3 missing fixes along the trajectory, 0.01% of the total path. At this point, AG195’s clean, regular, and complete trajectory (*n* = 26280) is ready for analysis with `stmove`.

### Individual Metrics

Running ‘build report’ with our complete, and regular trajectory for AG195, three plots are provided to begin: an *x*-*y* plot of the path (Fig. 4) and two histograms (Fig. 5) showing the distribution of step sizes (meters) and the distribution of turning angles (radians). Investigating these plots, we easily identify the range and most common step sizes of this elephant, based on a sampling interval of 20 mins: it most commonly moves less than 100 meters; but, on rare occasions, it moves upwards of a kilometer in this time interval. From the empirical distribution of turning angles we see that individual AG195 does not have a preferred turn direction and, from the strong peak in the histogram around 0 radians, some correlation is evident in directional persistence over periods that exceed 20 minutes. These empirical distributions are useful for identifying general behavioral profiles or outliers. Additionally, it is common to sample from these same empirical distributions when simulating movement tracks for future analyses.

In a `stmove` report that includes rolling statistics, three plots are provided of the running values for mean_dist, mean_ang, acf_dist, acf_ang, and ccf. Plots of mean dist and mean_ang are, by default, faceted by month to handled long term datasets with high resolution. The third plot displays smoothed conditional means splines across all rolled values of acf_dist, acf_ang, and ccf to give a high level view of patterns in these statistics across the entire temporal extent of the trajectory. Note in Fig. 6, the clear increase in mean step size in the months of February and March 2010, possibly the marker of increased movement during the start of the wet season. Diurnal interval statistics are plotted in similar fashion, with separate splines for morning versus evening intervals to illuminate differences between them. Coarser intervals, i.e. lunar or seasonal, are plotted using stair-step plots or bar charts to clearly demonstrate how estimates change from one interval to the next (Table 1).

**Table 1:** Interval Statistics. An example of Lunar statistical output (AG195)

| interval_start | phase | mean_dist | sd_dist | acf_dist | mean_ang | sd_ang | acf_ang | ccf |
| --- | --- | --- | --- | --- | --- | --- | --- | --- |
| [1,12) | Full-Waning | 99.827 | 125.683 | 0.571 | -0.001 | 1.313 | -0.035 | 0.028 |
| [12,26) | New-Waxing | 117.765 | 122.113 | 0.517 | 0.015 | 1.217 | -0.061 | -0.012 |
| [26,41) | Full-Waning | 180.998 | 238.705 | 0.686 | -0.036 | 1.317 | 0.020 | 0.041 |
| [41,56) | New-Waxing | 244.072 | 284.247 | 0.710 | 0.064 | 1.300 | 0.023 | -0.026 |
| [56,71) | Full-Waning | 210.297 | 253.647 | 0.674 | 0.065 | 1.366 | 0.043 | -0.018 |

When wavelet analyses are included within a report they can be applied to any of the 5 primary statistics and return individual power spectrum plots (Fig. 7). As an exploratory data analysis tool, these plots are intended to illuminate periodicity in the trajectory if it is present; for more information on wavelet transformations and the interpretation of these plots, we direct readers to Torrence and Combo’s practical guide to wavelet analysis [36]. Also, for examples of interpreting wavelet plots in movement ecology see [4, 5].

Finally, our full report for AG195 includes two methods of home range construction: a *k*-LoCoH hull set and an auto-correlated kernel density estimation both plotted with 25, 50 and 90% isopleths (Fig. 8). The methods differ profoundly and produce notably different estimations of space use with *k*-LoCoH often being quite restrictive and AKDE offering larger estimates with confidence bands. LoCoH methods provide clear information of where the animals have been and the areas locally bounded by the relocation points, while AKDE provides projections of where the animal is likely to be found if environmental and landscape features do not play a role in the animals movement behavior.

A final note of caution for users, although we have demonstrated a report including examples from all possible analyses here, which analyses are relevant to study at hand will be dependent on many variables including animal behavior, sampling rate and sampling duration—and, of course, questions of interest. Although animal movement is inherently a continuous process, relocation sampling is a discrete process in space and time. The sampling resolution influences the analyses and the conclusions one is able to draw regarding animal movement behavior [37]. Therefore, consideration of the resolution of your data is crucial before deciding which statistics to include in your custom report. Rolling and diurnal statistics, as well as wavelet plots, often are more appropriate for trajectories with higher resolution data. Interval statistics at the lunar or seasonal level are appropriate for larger-grain data, provided that sampling continued for long enough. Both home-range construction options can be used on data at any resolution, although AKDE is more appropriate at relatively high temporal resolution (on the order of minutes and fractions thereof) while LoCoH methods are appropriate and lower levels of temporal resolution (hours and large fractions thereof). We encourage users to think critically about the nature of their data before conducting even the most basic exploratory data analyses.

## 4 Conclusions

In 2008, Ran Nathan *et al.* [38] laid out the movement ecology paradigm, which effectively situated the emerging discipline within the broader ecological context, but fell short of dictating a set of baseline analyses that should be run on any newly-collected movement data. The movement ecology paradigm has informed the hypothesis generation process and guided the data collection procedures of innumerable studies, but the absence of a core set of standardized analyses among the many novel tools available to researchers has made it difficult to contextualize the movement patterns of an animal or species and to compare across studies and wildlife. `stmove` strives to fill the gap and make it easy for researchers to employ a standardized set of tools that provide basic insights into the movement of individuals. While this package is primarily an opinionated wrapper around other open-source contributors’ work and packages, the primary advantage and goal of this package is to provide a simple, single-command procedure to produce comprehensible and customized reports covering important baseline analyses one should conduct on GPS movement data. Once these analyses have been undertaken and properly interpreted, one can then pursue various kinds of analysis that address subsequent questions of interest (e.g., generalized linear models of location and landscape [39], hidden Markov modeling to identify behavioral states [40, 41], or step selection analyses [42, 43]). This package strives to set a standard for what is minimally needed before embarking on such analyses and in doing so hopes to provide a framework and infrastructure that democratizes foundational movement analysis and enhances comparability of studies.

## Notes

https://github.com/dpseidel/stmove

## References

[1] Joo, R., Boone, M.E., Clay, T.A., Patrick, S.C., Clusella-Trullas, S., Basille, M.: Navigating through the r packages for movement. arXiv preprint 1901.05935 (2019)

[2] R Core Team: R: A Language and Environment for Statistical Computing. R Foundation for Statistical Computing, Vienna, Austria (2018). R Foundation for Statistical Computing. https://www.R-project.org/

[3] Morales, J.M., Haydon, D.T., Frair, J., Holsinger, K.E., Fryxell, J.M.: Extracting more out of relocation data: building movement models as mixtures of random walks. Ecology 85(9), 2436–2445 (2004)

[4] Wittemyer, G., Polansky, L., Douglas-Hamilton, I., Getz, W.M.: Disentangling the effects of forage, social rank, and risk on movement autocorrelation of elephants using fourier and wavelet analyses. Proceedings of the National Academy of Sciences, 0801744105 (2008)

[5] Polansky, L., Wittemyer, G., Cross, P.C., Tambling, C.J., Getz, W.M.: From moonlight to movement and synchronized randomness: Fourier and wavelet analyses of animal location time series data. Ecology 91(5), 1506–1518 (2010)

[6] Getz, W.M., Wilmers, C.C.: A local nearest-neighbor convex-hull construction of home ranges and utilization distributions. Ecography 27(4), 489–505 (2004)

[7] Worton, B.: A review of models of home range for animal movement. Ecological modelling 38(3-4), 277–298 (1987)

[8] Calenge, C.: The package “adehabitat” for the r software: a tool for the analysis of space and habitat use by animals. Ecological modelling 197(3-4), 516–519 (2006)

[9] Moritz, S., Bartz-Beielstein, T.: imputeTS: Time Series Missing Value Imputation in R. The R Journal 9(1), 207–218 (2017)

[10] Patterson, T.A., Thomas, L., Wilcox, C., Ovaskainen, O., Matthiopoulos, J.: State–space models of individual animal movement. Trends in ecology & evolution 23(2), 87–94 (2008)

[11] Jonsen, I.D., Flemming, J.M., Myers, R.A.: Robust state–space modeling of animal movement data. Ecology 86(11), 2874–2880 (2005)

[12] Long, J.A.: Kinematic interpolation of movement data. International Journal of Geographical Information Science 30(5), 854–868 (2016)

[13] Bar-David, S., Bar-David, I., Cross, P.C., Ryan, S.J., Knechtel, C.U., Getz, W.M.: Methods for assessing movement path recursion with application to african buffalo in south africa. Ecology 90(9), 2467–2479 (2009)

[14] Berger-Tal, O., Bar-David, S.: Recursive movement patterns: review and synthesis across species. Ecosphere 6(9), 1–12 (2015)

[15] Abrahms, B., Seidel, D.P., Dougherty, E., Hazen, E.L., Bograd, S.J., Wilson, A.M., McNutt, J.W., Costa, D.P., Blake, S., Brashares, J.S., et al.: Suite of simple metrics reveals common movement syndromes across vertebrate taxa. Movement ecology 5(1), 12 (2017)

[16] Wickham, H.: Ggplot2: Elegant Graphics for Data Analysis. Springer, New York (2016). https://ggplot2.tidyverse.org

[17] Ushey, K.: RcppRoll: Efficient Rolling / Windowed Operations. (2018). R package version 0.3.0. https://CRAN.R-project.org/package=RcppRoll

[18] Ulrich, J.: TTR: Technical Trading Rules. (2018). R package version 0.23-4. https://CRAN.R-project.org/package=TTR

[19] Edelhoff, H., Signer, J., Balkenhol, N.: Path segmentation for beginners: an overview of current methods for detecting changes in animal movement patterns. Movement Ecology 4(1), 21 (2016)

[20] Seidel, D.P., Dougherty, E., Carlson, C., Getz, W.M.: Ecological metrics and methods for gps movement data. International Journal of Geographical Information Science 32(11), 2272–2293 (2018)

[21] Gurarie, E., Andrews, R.D., Laidre, K.L.: A novel method for identifying behavioural changes in animal movement data. Ecology letters 12(5), 395–408 (2009)

[22] Gurarie, E., Bracis, C., Delgado, M., Meckley, T.D., Kojola, I., Wagner, C.M.: What is the animal doing? tools for exploring behavioural structure in animal movements. Journal of Animal Ecology 85(1), 69–84 (2016)

[23] Lazaridis, E.: Lunar: Lunar Phase & Distance, Seasons and Other Environmental Factors. (2014). (Version 0.1-04). http://statistics.lazaridis.eu

[24] Bunn, A.G.: A dendrochronology program library in r (dplr). Dendrochronologia 26(2), 115–124 (2008). doi:10.1016/j.dendro.2008.01.002

[25] Bunn, A., Korpela, M., Biondi, F., Campelo, F., Mérian, P., Qeadan, F., Zang, C., Pucha-Cofrep, D., Wernicke, J.: dplR: Dendrochronology Program Library in R. (2018). R package version 1.6.9. https://CRAN.R-project.org/package=dplR

[26] Fieberg, J., Börger, L.: Could you please phrase “home range” as a question? Journal of Mammalogy 93(4), 890–902 (2012)

[27] Lyons, A., Getz, W., R Development Core Team: T-LoCoH: Time Local Convex Hull Homerange and Time Use Analysis. (2018). R package version 1.40.05

[28] Calabrese, J.M., Fleming, C.H., Gurarie, E.: ctmm: an r package for analyzing animal relocation data as a continuous-time stochastic process. Methods in Ecology and Evolution 7(9), 1124–1132 (2016)

[29] Fleming, C.H., Calabrese, J.M.: A new kernel density estimator for accurate home-range and species-range area estimation. Methods in Ecology and Evolution 8(5), 571–579 (2017)

[30] Noonan, M.J., Tucker, M.A., Fleming, C.H., Akre, T.S., Alberts, S.C., Ali, A.H., Altmann, J., Antunes, P.C., Belant, J.L., Beyer, D., et al.: A comprehensive analysis of autocorrelation and bias in home range estimation. Ecological Monographs 89(2), 01344 (2019)

[31] Getz, W.M., Fortmann-Roe, S., Cross, P.C., Lyons, A.J., Ryan, S.J., Wilmers, C.C.: Locoh: nonparameteric kernel methods for constructing home ranges and utilization distributions. PloS one 2(2), 207 (2007)

[32] Lyons, A.J., Turner, W.C., Getz, W.M.: Home range plus: a space-time characterization of movement over real landscapes. Movement Ecology 1(1), 2 (2013)

[33] Allaire, J., Xie, Y., McPherson, J., Luraschi, J., Ushey, K., Atkins, A., Wickham, H., Cheng, J., Chang, W., Iannone, R.: Rmarkdown: Dynamic Documents for R. (2018). R package version 1.11. https://rmarkdown.rstudio.com

[34] Xie, Y., Allaire, J.J., Grolemund, G.: R Markdown: The Definitive Guide. Chapman and Hall/CRC, Boca Raton, Florida (2018). ISBN 9781138359338. https://bookdown.org/yihui/rmarkdown

[35] Tsalyuk, M., Kilian, W., Reineking, B., Getz, W.M.: Temporal variation in resource selection of african elephants follows long-term variability in resource availability. Ecological Monographs, 01348 (2019)

[36] Torrence, C., Compo, G.P.: A practical guide to wavelet analysis. Bulletin of the American Meteorological Society 79(1), 61–78 (1998)

[37] Codling, E., Hill, N.: Sampling rate effects on measurements of correlated and biased random walks. Journal of Theoretical Biology 233(4), 573–588 (2005)

[38] Nathan, R., Getz, W.M., Revilla, E., Holyoak, M., Kadmon, R., Saltz, D., Smouse, P.E.: A movement ecology paradigm for unifying organismal movement research. Proceedings of the National Academy of Sciences 105(49), 19052–19059 (2008)

[39] Manly, B., McDonald, L., Thomas, D.L., McDonald, T.L., Erickson, W.P.: Resource Selection by Animals: Statistical Design and Analysis for Field Studies. Springer, ??? (2007)

[40] Patterson, T.A., Basson, M., Bravington, M.V., Gunn, J.S.: Classifying movement behaviour in relation to environmental conditions using hidden markov models. Journal of Animal Ecology 78(6), 1113–1123 (2009)

[41] Michelot, T., Langrock, R., Patterson, T.A.: movehmm: an r package for the statistical modelling of animal movement data using hidden markov models. Methods in Ecology and Evolution 7(11), 1308–1315 (2016)

[42] Thurfjell, H., Ciuti, S., Boyce, M.S.: Applications of step-selection functions in ecology and conservation. Movement ecology 2(1), 4 (2014)

[43] Avgar, T., Potts, J.R., Lewis, M.A., Boyce, M.S.: Integrated step selection analysis: bridging the gap between resource selection and animal movement. Methods in Ecology and Evolution 7(5), 619–630 (2016)

